# Spatial learning overshadows learning novel odors and sounds in both a predatory and a frugivorous bat

**DOI:** 10.1101/2022.06.03.494735

**Authors:** M. May Dixon, Gerald Carter, Michael J. Ryan, Rachel A. Page

## Abstract

To be efficient while foraging, animals should be selective about attending to and remembering the cues of food that best predict future meals. One hypothesis is that animals with different foraging strategies should vary in their reliance on spatial and feature cues. Animals that store food or feed on spatially stable food, like fruit or flowers, should be predisposed to learning a meal’s location, whereas predators that hunt mobile prey should instead be biased towards learning feature cues such as color or sound. Previous studies suggest that nectar- and fruit-feeding bats should rely relatively more on spatial cues, whereas predatory bats should rely more on feature cues, yet no experiment has compared these two foraging strategies under the same conditions. To test this hypothesis, we compared learning in the frugivorous bat, *Artibeus jamaicensis*, and the predatory bat, *Lophostoma silvicolum*, which hunts katydids using acoustic cues. We trained bats to find food paired with a unique odor, sound, and location. We then dissociated these cues to assess which cue each bat had learned. Rather than finding that the frugivore and predator are on opposite ends of a continuum in their relative reliance on spatial and feature cues, we found that both species learned spatial cues with no evidence that either learned the sounds or odors. We discuss interpretations of these results in the context of past work on use of spatial cues versus feature cues. Compared to feature cues, spatial cues may be fundamentally more rich, salient, or memorable.

**Lay Summary:** When animals learn how to obtain food, they could attend to many possible cues—such as the associated smells, sights, sounds, or locations of the food. It has been hypothesized that natural selection has shaped animals to pay particular attention to the types of cues that are most useful for finding their typical food. In bats, this would predict that predatory species that hunt mobile, noisy insects by sound should be biased towards learning a food-associated sound, whereas bats that eat stationary and silent fruit should be biased towards learning a food’s location. We gave predatory and fruit-eating bats the opportunity to simultaneously learn food-rewarded odors, sounds, or locations.

Regardless of their diets, both species of bats learned locations rather than the odors or sounds associated with food. Spatial cues may be particularly rich or salient, and when they are reliable, animals may use them for learning over other cues regardless of their typical food.

## Introduction

Diet shapes the evolution of almost every aspect of an animal’s biology from morphology to sensory and cognitive systems (MacLean et al. 2012; Stevens 2014; Rosati 2017; Amodio et al. 2019). If an animal can reliably find or evaluate food using a particular sensory modality, such as smell, there may be selection to be more sensitive in that modality (Warrant 2016) and but also to attend to and learn odor-food associations more easily than associations with cues in other sensory modalities (Garcia and Koelling 1966; Dunlap and Stephens 2014). For example, fruit flies in the lab can evolve to learn odor associations better than color associations when odor is a more reliable indicator of a safe place to lay eggs (Dunlap and Stephens 2014).

Strong evidence that natural selection shapes the cues animals learn comes from animals that need to remember the location of hidden food. Scatter-hoarding species rely more on spatial cues (e.g., absolute position in space) than object-specific cues (e.g., shape or color) as compared to related species that do not scatter-hoard (Barkley & Jacobs, 2007; Pravosudov & Roth II, 2013; Sherry et al., 1992; Shettleworth, 2003; Table S1). For example, when black-capped chickadees, *Poecile atricapillus*, were trained to find food that was simultaneously associated with a color pattern, a position in an array, and an absolute location in a room, they preferentially relied on absolute location, whereas non-caching dark-eyed juncos, *Junco hyemalis*, relied on the three cues equally (Brodbeck 1994). Further experiments showed that this difference in cue use was a result of a greater ability to remember spatial cues, rather than a difference in attention or spatial discrimination ability (Shettleworth 2003). Presumably, absolute location is the most reliable cue for refinding hidden food after time has passed and other cues in the landscape might have changed.

Location may also be the most reliable cue for animals that frequently return to a renewable resource. The rufous hummingbird, *Selasphorus rufus*, repeatedly visits flowers in its territory throughout the day and must remember their locations and states. To refind flowers in experiments, it preferentially relied on spatial cues (both position relative to other flowers and absolute location) over feature cues (color and pattern) (Healy & Hurly, 1998; Hurly & Healy, 1996; Table S1).

While many studies show that animals that forage on spatially predictable food prefer using spatial cues (Table S1), there has been less attention to the other side of this continuum: animals for which spatial cues of food are unreliable, but feature cues are informative. For many predators that consume ephemeral or mobile prey, feature cues are more predictive of a future meal than spatial cues. Yet, there are comparatively few examples of animals preferring feature cues over spatial cues; these include studies of European greenfinches (*Chloris chloris*) (Herborn et al. 2011), domestic chicks (*Gallus gallus*) (Vallortigara 1996), and humans (*Homo sapiens*) (Haun et al. 2006), and in each of these studies, only some treatments led to a preference for spatial cues. European greenfinches, which eat ephemeral berries, chose color over spatial cues but only when exposed to the rewarded combination of a color and location once; they switched to spatial cues when trained with the compound cue 10 times (Herborn et al. 2011). Another experiment found that female chicks preferred color cues whereas males preferred spatial cues (Vallortigara 1996). Among human children, 3-year-olds relied on feature cues over spatial cues, but 1-year-olds relied on spatial cues, as did orangutans, gorillas, bonobos, and chimpanzees (Haun et al. 2006). The relative salience of spatial and feature cues is likely to be contextual—depending on the qualities of the cues offered (LaDage et al. 2009), but it is interesting that there are relatively few examples of animals exclusively preferring to use feature cues when offered spatial cues. In several cases, animals predicted to rely on feature cues have instead relied on spatial cues when both cues were available (Williams 1967a; Williams 1967b; Hodgson and Healy 2005; Daneri et al. 2011).

Since prey tend to be mobile, predators learning about a new prey species are expected to rely more on the feature cues of a new food, such as how the food sounds, looks, or smells, over the location where it was encountered. However, there is surprisingly little support for this preference in the literature on controlled tests comparing learning of spatial and feature cues of food; to our knowledge the study on female chicks is the only case (Vallortigara 1996).

Stich and Winter (2006) proposed that species of New World leaf-nosed bats (Phyllostomidae) should vary on a continuum in their reliance on spatial versus feature cues based on the extent to which their food is spatially stable (‘niche-specific cognitive strategies’). On one end of the spectrum, nectar bats, like hummingbirds, were hypothesized to rely relatively more on spatial cues because they have to remember and refind multiple flowers dispersed in kilometers of jungle (Stich and Winter 2006). Frugivorous bats were hypothesized to show intermediate behaviors— relying more on spatial cues because these bats could benefit from remembering the locations of profitable trees, but once at the tree, relying on odor to detect ripe fruit. On the other end of the spectrum, predatory bats were predicted to rely relatively less on spatial associations because their food is spatially unpredictable, and relatively more on feature cues such as prey shapes and sounds (Stich and Winter 2006; Hulgard and Ratcliffe 2014).

The hypothesis of niche-specific cognitive strategies has been supported by studies on species that should rely more on spatial cues (Thiele and Winter 2005; Carter et al. 2010; Henry and Stoner 2011). In captivity, the nectar bats in the genus *Glossophaga* learned spatial positions faster than feature associations (Stich and Winter 2006; Carter et al. 2010). *Glossophaga commissarisi* preferentially relied on spatial cues over feature cues (Thiele and Winter 2005). *Glossophaga soricina*, which sometimes eats insects, was outperformed by a more specialized nectivore, *Leptonycteris yerbabuenae*, on a spatial memory task (Henry and Stoner 2011), presumably because the specialist faces greater selection for remembering flower locations. A test comparing *G. soricina* and the frugivorous *Carollia perspicillata* found that both species relied on spatial cues and showed little evidence of learning odor or shape cues, despite all three cues being available (Carter et al. 2010).

A problem, however, is that all these species are predicted to rely heavily on spatial cues; the hypothesis has not been tested using bats that are predicted to rely strongly on feature cues. The logical next step is to compare use of spatial and feature cues in bat species with different predictions. Here, we compare cue selection in a frugivorous and a predatory phyllostomid bat. We chose the fruit-eating bat species, *Artibeus jamaicensis*, because it specializes on figs (Ortega and Castro-Arellano 2001) and finds them primarily using scent (Kalko et al. 1996), and we chose the predatory bat, *Lophostoma silvicolum*, because it finds katydids by listening to their mating calls (Tuttle et al. 1985; Belwood 1988; Falk et al. 2015). Notably, these acoustic eavesdroppers are expected to rely on feature cues even at a distance, because they eavesdrop on the sexual advertisement calls of their prey from a large distance (Page et al. 2012; Jordan and Ryan 2015). To test which cues the bats had associated with the food, we first allowed the subjects to learn to forage at one rewarded feeder in an array of feeders, each with its own unique combination of odor, sound, and location. Next, we separated two of the rewarded cues (such as sound and location) and removed the third (such as odor), and then observed which feeder the bats chose. If the spatial stability of their food predicts how much a species relies on spatial vs feature cues, then we predicted that the insectivore would rely relatively more on sound cues and that the frugivore would rely relatively more on spatial cues and odor cues.

## Methods

### Capture and care

We trained and tested 12 male *Artibeus jamaicensis* and 10 male *Lophostoma silvicolum* caught from Soberanía National Park, Panamá and the surrounding forest, in either mist nets at night or in their roosts during the day. *Artibeus jamaicensis* were trained in two cohorts of six bats each, and *L. silvicolum* were trained in four cohorts of two or three bats. Bats were maintained in a large open-air flight room (5 m × 5 m × 2.5 m) with a cloth roost in the corner, except on the night of capture and briefly during the nights of testing, when they were kept in small tents (∼ 1.2 m × 0.75 m × 1 m). Water was provided *ad libitum* from trays on the floor. *A. jamaicensis* were fed a mix of banana, papaya, and melon, and *L. silvicolum* ate thawed katydids. At the end of the experiment, all bats were injected with passive integrated transponders (PIT tags) to prevent re-testing and returned to the wild.

### Experimental apparatus and stimulus generation

Food was presented to bats on wood platform-feeders that were 39.5 cm × 29 cm and 90 cm tall, with holes in the top to allow odor and sound cues to come through from a compartment below (Figure 1). The four odor cues were ultra-concentrated candy oils (cinnamon, anise, almond, and sassafras; LorAnn oils, (O’Mara et al. 2014)), placed in 1.5 ml vials with a cotton wick. The four sound cues were created from four different cell phone ringtones that were modified to have peaks in frequencies at 5, 18, and 20 kHz (Figure S1), near the peak hearing sensitivities for *A. jamaicensis* (Heffner et al. 2003) and *L. silvicolum* (Geipel et al. 2021). The peak amplitude was normalized, and the sounds were broadcast at 65 dB SPL at 1 m (re 20 µPa) through Fostex speakers (P650k) via Pyle PCA2 stereo power amplifiers. We knew that all sound cues were audible to the bats because in pilot studies, both species oriented their bodies towards the speaker when each of the sound cues was played at the experimental amplitudes.

**Figure 1:**
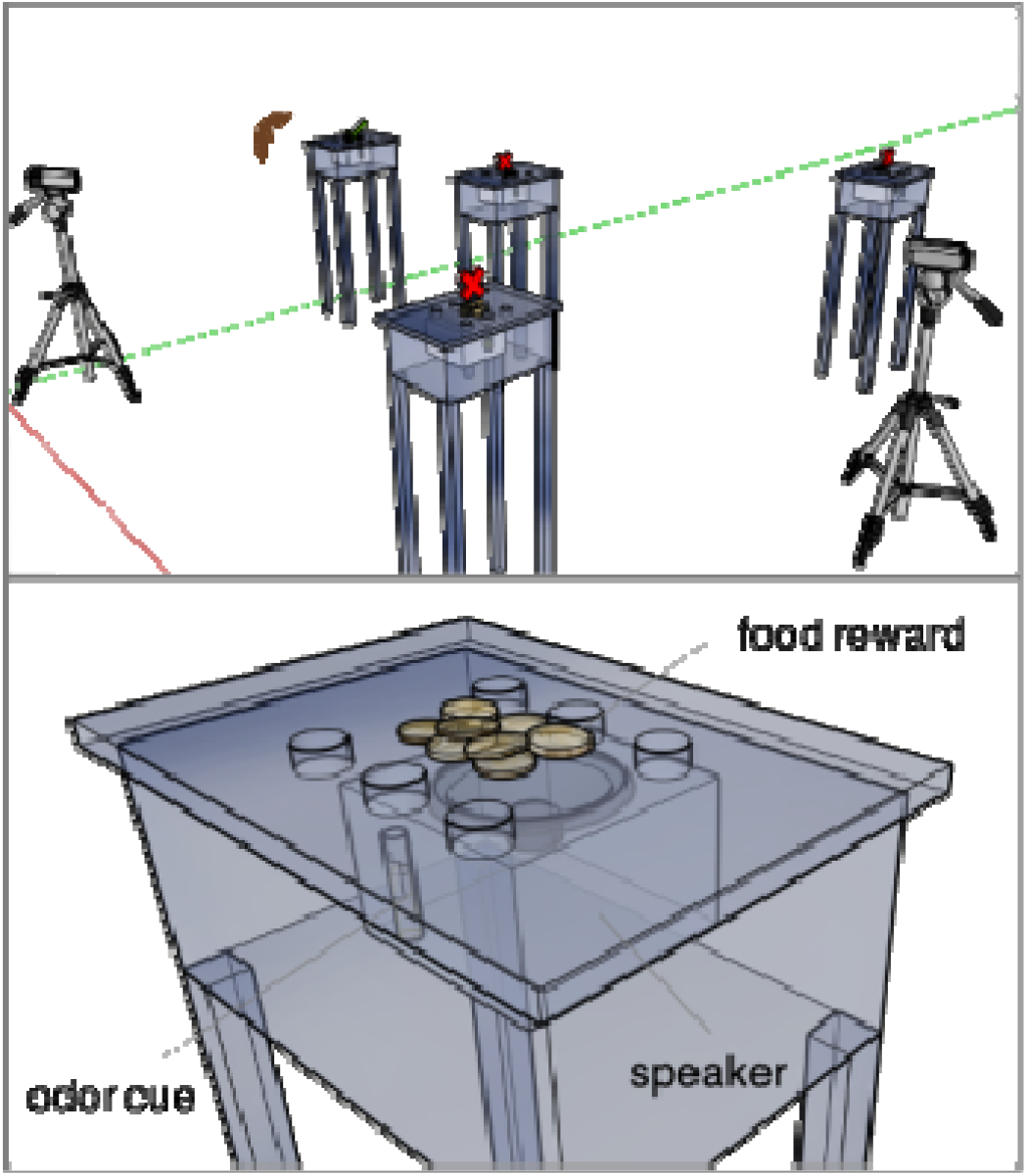
Diagram of the experimental feeders and set-up. Top: Experimental set-up. Feeders were positioned in unique locations in the flight cage, and food was placed on top. Only one feeder was rewarded (indicated by green check), while the food on the other feeders was rendered inedible with ascorbic acid (indicated by red crosses). Food was placed on top. Bottom: Experimental feeder. Feeder platforms were each equipped with a speaker that played a unique sound and a vial containing fragrant candy oil. Feeders were provisioned with either pieces of banana (pictured), or katydids.

To create unique spatial locations, we made a 4 m × 4 m grid on the floor and used a random number generator to assign the feeders to coordinates with the condition that feeders could not be within a meter of one another. Sounds and odors were also randomly assigned to each feeder. The flight room was illuminated with a 25-W red light bulb and at least four infrared lights (Clover Electronics IR045). Tests were filmed with two Sony Handycam DCR-SR45 cameras placed in corners of the flight room.

### Pre-training

Bats were trained to retrieve food off the target feeder. First, on the night of capture, bats were fed by hand inside a small tent to acclimate them to captivity. On the second night, they were trained to eat food from the feeders (without the experimental cues) inside the tent. We placed both fruit and katydids on the speakers during this phase and played a *Eubliastes pollonerae* katydid call at 15 s intervals at 65 dB SPL until all bats retrieved food from the feeder. To further entice *L. silvicolum* to the feeder, we played calling songs of different katydid species known to elicit approaches (e.g., *Docidocercus gigliotosi, Anapolisia colosseum*, (Falk et al. 2015)) until all bats retrieved food.

### Training

When bats were flying consistently to the feeders, we released them into the larger flight room. To elicit successful feeding, on the first night in the flight room we only set out the rewarded target feeder, in the rewarded location with the rewarded odor and rewarded sound cue. This set-up was sufficient to elicit feeding visits from all the *A. jamaicensis*. For some of the *L. silvicolum*, we had to additionally broadcast low amplitude calling songs of the aforementioned katydid species to coax the bats to the target feeder initially.

The following night, we placed all four feeders in the room, each with their respective sounds, odors, and locations (with no additional acoustic cues). To control for the effect of the food scent, we placed food on top of all feeders but rendered the food on the non-rewarded feeders inedible by applying ascorbic acid (vitamin C) to its surface (1.25 g per 20 g of food). In pilot tests we determined that this concentration of ascorbic acid, though harmless, is sour and aversive to the bats. We periodically replenished the food to ensure food was present at all feeders. To allow the bats to learn which feeder was rewarded, they foraged ad libitum in this set-up for one week.

### Cue-learning tests

After pre-training, all bats were moved to a holding tent in an adjacent room and each bat was tested individually in the flight room. This test was designed to ensure that the bats had learned the association between the target feeder and the reward, and were not finding the target feeder using social cues or cues from the ascorbic acid. Tests lasted one hour. The set-up was the same as in training except that all the feeders were rewarded (no ascorbic acid). To proceed to the cue-dissociation tests, bats had to fly to the target feeder at least 5 times and make no more than two incorrect choices (probability of passing by chance is ∼1%). If the bat did not pass the cue-learning test, it was tested again on the following night. After all tests were run each evening, all bats were returned to the flight room with the pre-training set-up to eat to satiation and to reinforce the association with the target feeder.

### Cue dissociation tests

When a bat passed the cue-learning test, it started cue dissociation testing on the following night. These tests were designed to probe how the bats had learned to find the food by putting the previously rewarded cues into conflict (Brodbeck 1994). Following Carter et al. (2010), we ran tests with two of the three cues present simultaneously, thus allowing us to determine if each cue was learned and its relevant salience to the bat. Each bat experienced all three cue combinations (location vs sound, location vs odor, and sound vs odor) one time, in random order, with one test per night for three consecutive nights. In each test, one of the cue types was removed entirely (e.g., all sound cues) and the remaining cues were switched between the remaining three feeders, so that the previously rewarded cues were at different feeders (Figure 2). None of the feeders were rewarded: they all had 10 pieces of food (bananas or katydids) rendered unpalatable by coating in ascorbic acid. This encouraged the bats to switch strategies to increase our likelihood of detecting if they had learned other rewarded cues. All tests ran for one hour and were recorded with video cameras placed in the corners of the flight room.

**Figure 2:**
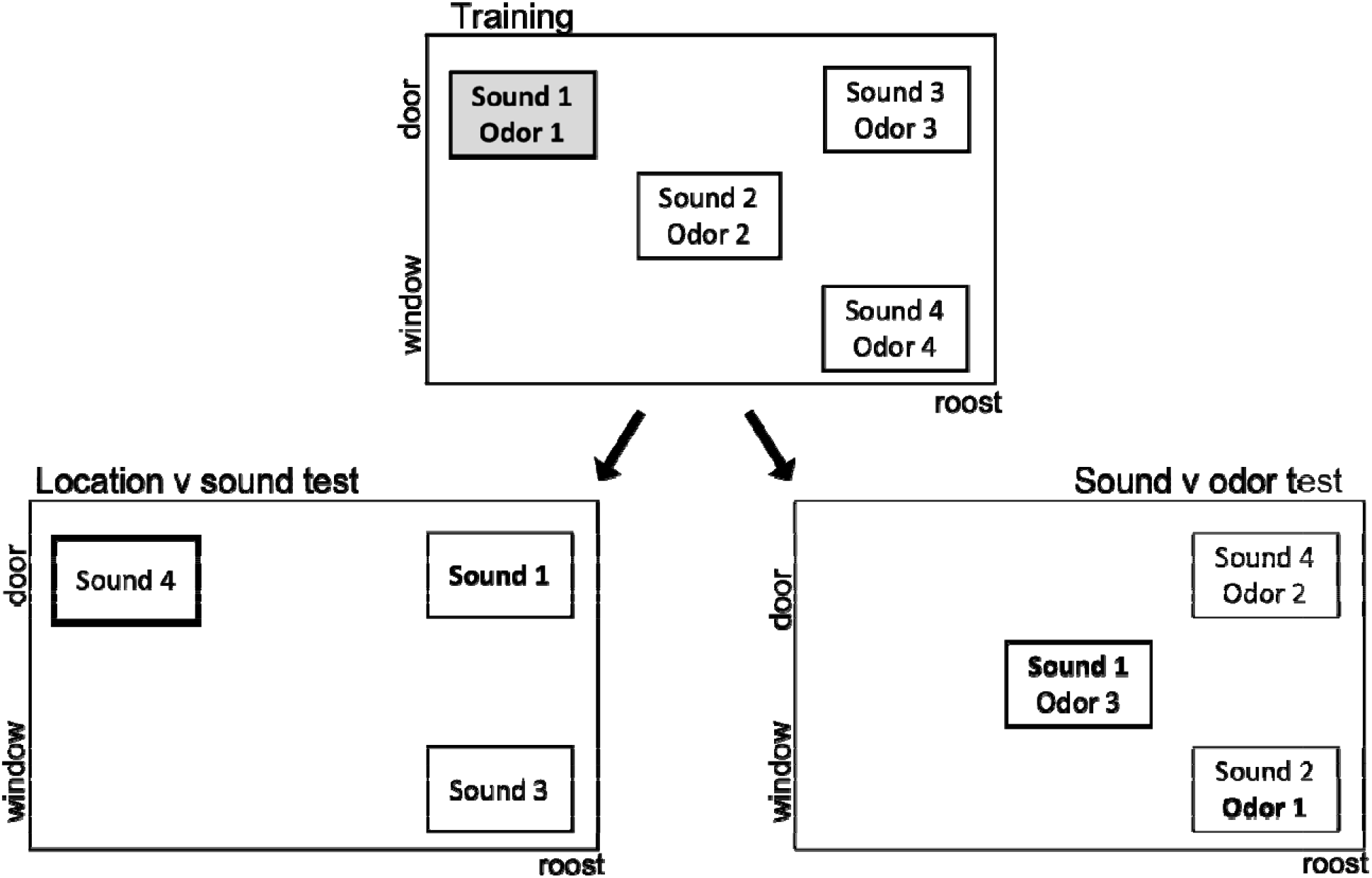
Experimental setup. Panels show examples of how cues were arranged in the flight cage, as seen from above, during training and cue-dissociation tests. Small rectangles represent the feeders, each with their own unique sound, odor, and location in the room. During training (top) one feeder was rewarded (shaded rectangle, bold font and heavy border), and the food in the other feeders was distasteful (open rectangles). During testing (bottom) no feeders were rewarded, and bats could choose between three feeders: two of which had one previously rewarded cue (bold font and heavy border), and one which had no previously rewarded cues. In the location versus sound test (bottom left), there were no odor cues, the sound cues were randomly swapped between feeders, and one feeder was removed. In the sound versus odor test (bottom right), the feeder at the rewarded location was removed, and the sounds and odors were swapped between the remaining feeders. Figure not to scale.

#### Location versus odor tests

In location versus odor tests, all the sounds were removed along with one of the non-rewarded feeders. The three remaining odors were switched at random but in a way that the previously rewarded odor was in a new position. Bats then chose between the previously rewarded location with an unrewarded odor, the previously rewarded odor in an unrewarded location, and a control feeder with a location and odor that had never been rewarded.

#### Location versus sound tests

In location versus sound tests, all of the odors were removed along with one of the three non-rewarded feeders. The three remaining sound stimuli were switched at random such that the previously rewarded sound was in a new position. Bats then chose between the feeder in the previously rewarded location broadcasting a previously unrewarded sound, the feeder with the previously rewarded sound in a previously unrewarded location, and a control feeder with a location and sound that had never been rewarded.

#### Sound versus odor tests

In sound versus odor tests, the feeder at the previously rewarded location was removed, along with a random non-rewarded odor and sound. The six remaining sounds and odors were switched at random such that the previously rewarded odor and sound were at different, previously unrewarded feeders. Bats then chose between the previously rewarded sound, the previously rewarded odor, and a control feeder with a sound, odor, and location that had never been rewarded.

### Behavioral analysis

To analyze the bats’ behavior, we video-recorded the feeders to score which one a bat chose first and the total number of choices to each feeder. Choices were counted if the bat hovered within 1 body length (about 7 cm) from the top of a feeder for more than 0.1 s (3 video frames) or if they touched down atop a feeder.

### Statistical analysis

To determine whether bats of each species first chose any of the three cues in a test more than expected by chance (33%), we first used an exact multinomial test. If a deviation from chance was detected, we used a binomial test to test if the most preferred cue was selected more than expected by chance.

As a measure of preference for each cue, we took the number of choices to each cue and subtracted the mean number of choices (i.e., the difference between the observed and expected value). We then used nonparametric bootstrapping (5000 permutations) to get the 95% confidence interval around the mean preference for each cue by species.

To test whether *A. jamaicensis* and *L. silvicolum* differed in their preferences for the three cues in each test, we used a permutation test. To do this, we first calculated the mean number of choices to each cue across all trials and calculated the species difference (*S*) as the mean choices by *A. jamaicensis* – mean choices by *L. silvicolum*. To create a null distribution of species differences (*S*) under the null hypothesis of random choices, we randomized the number of choices to each cue within each bat and test, and calculated *S*, repeating this 5000 times. Note that these permutations do not change the total numbers of choices per test and species, only the distribution of the choices within each test, so a difference between the observed and expected values are due to nonrandom choices by the bats. To get two-tailed p-values, we centered the distribution of expected values and calculated the proportion of absolute expected values greater than the absolute observed value. To get two-tailed p-values, we calculated the proportion of expected values that had a greater or equal difference from the mean of the expected distribution than the observed value. To get two-tailed p-values, we found the absolute difference between the mean of the expected distribution and the observed value and calculated the proportion of expected values that had a greater or equal absolute difference.

## Results

### Artibeus jamaicensis

#### Location versus odor tests

In tests where bats could choose between the previously rewarded location, the previously rewarded odor, and a control odor-location combination that had never been rewarded, *A. jamaicensis* did not make first choices randomly (α = 0.05, *N =* 9, *P =* 0.014; Figure 3a); seven of nine bats chose the location feeder first (bats expected by chance = 3, *P =* 0.008). Over the trials *A. jamaicensis* repeatedly chose the location feeder more often than odor and the control (Figure 3b).

**Figure 3:**
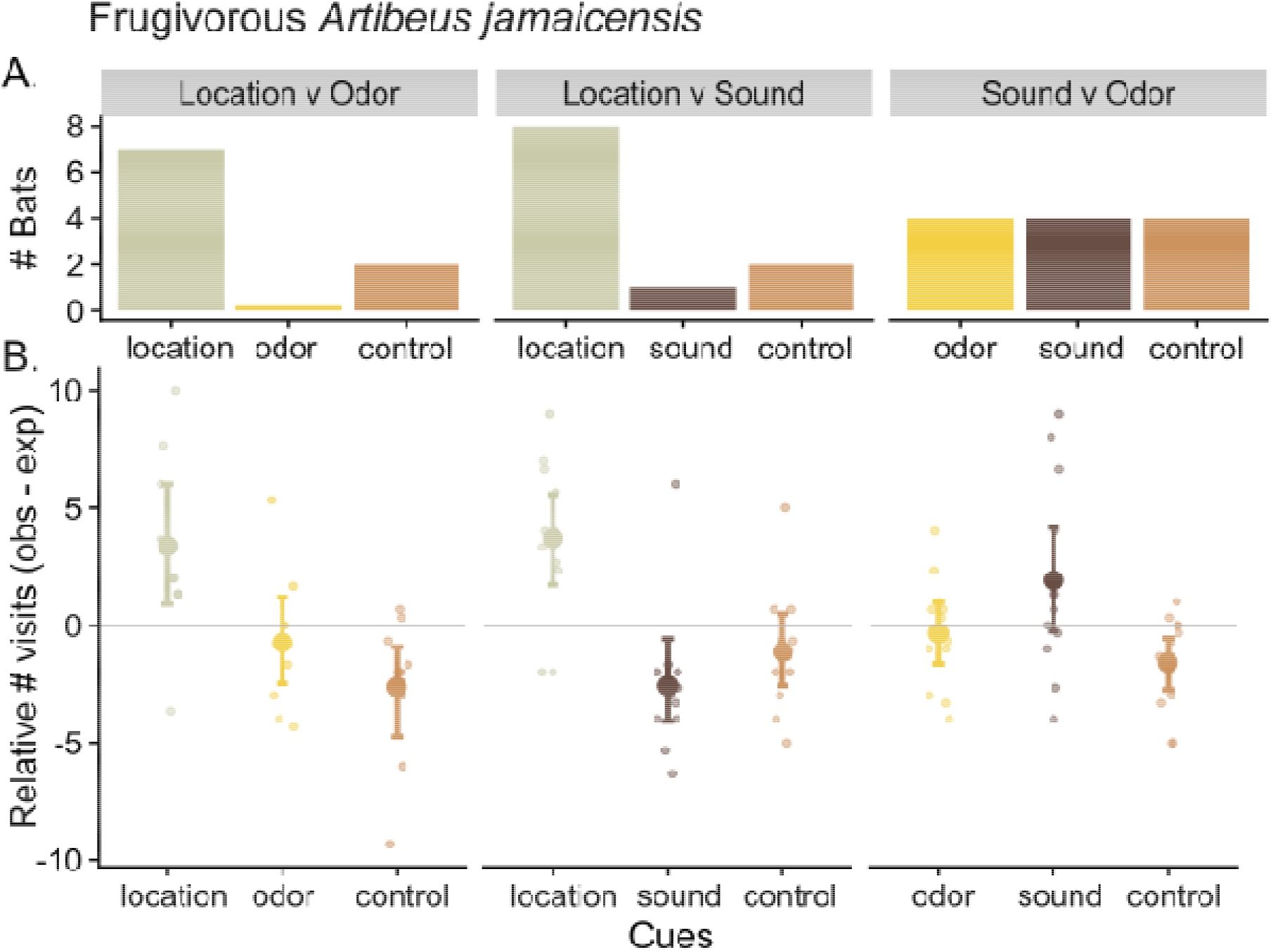
*Artibeus jamaicensis* choices to the cues in each of the three test types. A) The first choice of cue combination. B) The relative number of choices (choices to cue – mean choices) that bats made to each cue over one hour. Small points represent individual bats, large points and whiskers represent means and 95% confidence intervals.

#### Location versus sound tests

In tests where bats could choose the rewarded location, the rewarded sound, and a control, a location and sound that had never been rewarded, eight of 11 *A. jamaicensis* first chose location (bats expected by chance = 3.7, *P =* 0.009), one chose sound, and two chose the control (α = 0.05, *P =* 0.053; Figure 3a). Over the trials *A. jamaicensis* repeatedly chose location more often than sound and the control (Figure 3b).

#### Sound versus odor tests

In tests where bats could choose between the previously rewarded sound, the rewarded odor, and a control sound and odor combination that had never been rewarded, *A. jamaicensis* first choices were equal across the cues and consistent with random choice (α = 0.05, *N =* 12, *P >* 0.9; Figure 3b). Over the trials *A. jamaicensis* did not repeatedly choose any feeder more often than expected by chance, although they tended to choose sound more often than the control (Figure 3b).

#### Lophostoma silvicolum

##### Location versus odor tests

The predatory bat, *L. silvicolum*, chose some cues first more than expected by chance (α = 0.05, *N =* 10, *P =* 0.006; Figure 4a). Eight of 10 bats chose location first (3.3 expected by chance, *P =* 0.003), zero chose odor and two chose the control feeder. Over the trials *L. silvicolum* repeatedly chose location relatively more often than odor and the control (Figure 4b).

**Figure 4:**
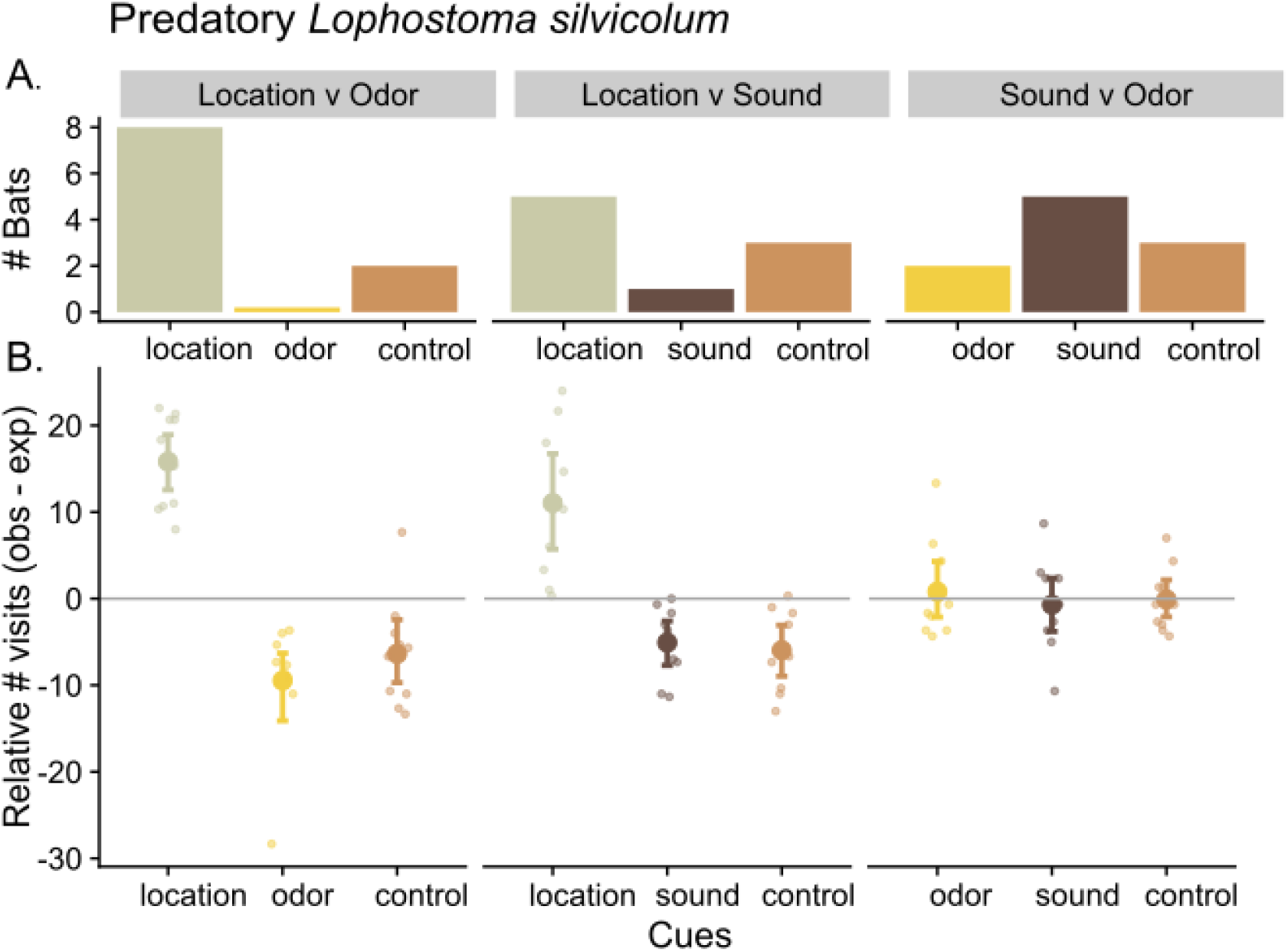
*Lophostoma silvicolum* choices to the cues in each of the three test types. A) The first choice of cue combination. B) The relative number of choices (choices to cue – mean choices) that bats made to each cue over one hour. Small points represent individual bats, large points and whiskers represent means and 95% confidence intervals.

##### Location versus sound tests

First choices by *L. silvicolum* did not deviate from random chance (α = 0.05, *N =* 9, *P* > 0.3; Figure 4a): five of eight chose location first, one sound, and three the control, but *L. silvicolum* repeatedly chose location significantly more often than it chose either sound or the control over the full trials (Figure 4b).

##### Sound versus odor tests

*L. silvicolum* first choices did not deviate from random chance (α = 0.05, *N =* 10, *P >* 0.6; Figure 4a). Five flew first to sound, two to odor, and three to the control^-^ feeder. Over the trials *L. silvicolum* did not repeatedly choose the sound feeders more often than the others (Figure 4b).

### Species differences

In location versus odor tests, *L. silvicolum* chose location relatively more often than *A. jamaicensis* (mean proportion of visits = 82%. vs 57% respectively; *S =* −19, *P =* 0.0014; Figure S2a), and odor relatively less often than *A. jamaicensis* (mean proportion of visits = 7% vs 25%; *S =* 2, *P =* 0.041; Figure S2b), but had similar proportion of choices to the control feeder (mean proportion of visits = 11% vs 18%; *S =* −3, *P >* 0.3; Figure S2c).

In location versus sound tests, *L. silvicolum* chose location relatively more often than *A. jamaicensis* (mean proportion of visits = 81% vs 60%; *S* = −8, *P =* 0.029; Figure S2d), but did not clearly choose the sound or control feeders more or less often (sound: mean proportion of choices = 10% vs 16%; *S =* 2, *P >* 0.4; Figure S2e; control: mean proportion of visits = 9% vs 24%; *S =* 4, *P >* 0.1; Figure S2f).

In sound versus odor tests, we saw no clear differences between species in the relative amount that they chose the three feeder types (mean proportion visits of visits were between 22 to 41%, *S =* −3 to 1, *P* > 0.1 for each, Figure S2g-i).

## Discussion

In this experiment we tested whether two bats with different foraging strategies would rely on cues differently when learning about a novel food item. If foraging strategy biases which cues are salient when learning about food, we predicted that the acoustic eavesdropping predator *L. silvicolum* would rely relatively more on sound cues than spatial cues, compared to the frugivorous *A. jamaicensis*, which is predicted to rely relatively more on spatial cues, followed by odor cues. However, we did not detect this pattern. Instead, both species used spatial cues more than the feature cues of sounds and odor to refind the novel food (Figure 3 and 4). Our results were consistent with spatial memory overshadowing the learning of novel sounds and odors. Indeed, there was no clear evidence that either species relied on the odor or the sound cues, even when the rewarded location was unavailable. If the two species did differ, the predatory *L. silvicolum* appeared to rely on spatial cues more than the frugivore, because it chose the location feeder relatively more often (Figure S2). This result does not support the hypothesis that predatory bats are cognitively specialized to rely relatively less on spatial cues than frugivorous bats while foraging (Stich and Winter 2006; Hulgard and Ratcliffe 2014). There are two main interpretations of this result.

First, phyllostomid bat species might not have clear diet-based cognitive specializations. All phyllostomid bats might flexibly shift their use of different cue types as the context changes, or they might always rely first on spatial cues. Given that they diversified relatively recently, ∼18-25 MYA (Monteiro and Nogueira 2011; Baker et al. 2012), it is possible that cognitive specializations might require more evolutionary time than the morphological divergences that are obvious in this lineage (e.g., Santana and Cheung 2016; Arbour et al. 2019).

Alternatively, subtle between-species differences in cue preference might exist yet be overshadowed by an overwhelming preference for use of spatial cues. The ability for spatial cues to overshadow feature cues has now been seen in all five bat species that have been tested in this paradigm (Table 1). Given that each of these species learns various feature cues (odors, echoacoustic images, or sounds) when foraging for food (Lemke 1984; Belwood 1988; Kalko et al. 1996; Thies et al. 1998; Patriquin et al. 2018; Brokaw et al. 2021), why might learning based on spatial cues appear so dominant as tested in this paradigm?

**Table 1:** Bat species tested in cue-dissociation experiments.

| **Guild** | **Species** | **Cues tested** | **Primary cue used** | **Secondary cue?** | **Citation** |
| --- | --- | --- | --- | --- | --- |
| Nectivore | *Glossophaga commissarisi* | Location (absolute), location (relative position), shape | Location (absolute) | Shape | (Thiele and Winter 2005) |
|  | *Glossophaga soricina* | Location, odor, shape | Location | Shape/ odor possibly | (Carter et al. 2010) |
| Frugivore | *Carollia perspicillata* | Location, odor, shape | Location | Shape/ odor possibly | (Carter et al. 2010) |
|  | *Artibeus jamaicensis* | Location, odor, sound | Location | None detected | Current study |
| Insectivore | *Lophostoma silvicolum* | Location, odor, sound | Location | None detected | Current study |

There are several non-mutually exclusive factors to consider. First, spatial learning may be fundamentally different from learning to associate specific sensory cues with a target object. Unlike odor and sound, the spatial cues in our study could be perceived in multiple senses (e.g., odor, vision, and echolocation), and represented in various ways (through egocentric, geometric, or landmark cues). ‘Locations’ are not single ‘cues’ but rather a collection of many possible cues, and as such may be more salient than any one type of cue. One possible implication is that, regardless of ecology, animals may have a bias towards using spatial cues whenever they are reliable (Day et al. 2003). Our study and several others used repeated training trials with a rewarded location (e.g., Williams 1967a; Williams 1967b; Hodgson and Healy 2005; Carter et al. 2010; Herborn et al. 2011), and this experimental design switched European greenfinches from relying on feature cues to spatial cues (Herborn et al. 2011). Therefore, animals that do not otherwise rely on spatial cues may flexibly learn to use them when they experience that a location is reliable.

A second related idea is that cue selection can depend on perceptual salience. For example, mountain chickadees normally rely primarily on spatial cues when finding food, but relied first on visual cues and secondarily on spatial cues when the visual task was much easier (two colors vs sixteen closely spaced locations) (LaDage et al. 2009). Nectar bats switched from using both flower shape and odor to using only odor cues when the foraging background was more complex (Muchhala and Serrano 2015). In our study, it is possible that discriminating the four locations was easier for the bats than discriminating the four sounds and odors used in the study.

Third, the assumption that frugivores and nectivores rely on spatial cues more than predatory bats in foraging could also be wrong, if for example, predatory bats use spatial memory extensively to return to previously profitable prey patches or hunting perches (Ratcliffe, 2009). For example, harbor seals use spatial memory to remember hunting grounds (Iorio-Merlo et al. 2022), and another predatory bat, *Megaderma lyra*, appears to use spatial memory to assess familiar hunting grounds which may reduce the need to echolocate when hunting (Ratcliffe et al. 2005).

Fourth, spatial learning is also critically important in these animals’ lives in contexts beyond learning about food. All the species in Table 1 forage in the rainforest interior, navigating nightly through dense cluttered jungle to find food and then return to their roosts. There is some evidence that foraging in cluttered space versus open space predicts bat spatial cognition (Clarin et al. 2013) and brain size (Safi and Dechmann 2005; Dechmann and Safi 2009). Selection for reliance on a strong spatial memory for homing or navigation may generalize to learning about food, such that any of these species will preferentially rely on a reliable spatial association when it is available, even when spatial associations are not good predictors of food in the wild. If so, foraging habitat may be a better predictor of cue reliance than foraging guild in bats (Odling-Smee and Braithwaite 2003; Cheng et al. 2014). This hypothesis could also help explain why other animals that are not predicted to rely extensively on spatial cues often prefer them (Williams 1967a; Williams 1967b; Hodgson and Healy 2005; Carter et al. 2010). It would be interesting to compare spatial and feature learning between closely related bat species that live in complex versus simple environments. If the challenge of navigating more cluttered environments selects for greater reliance on learning spatial cues, cue-reliance should vary across species that forage in the forest interior vs open space.

Finally, bats might have cognitive specializations that take a different form than the one we tested. We considered differences in how bats choose which novel cues to associate with food, but specializations may occur at other stages of cognition. For example, species have different ‘sensory filters’ that constrain what they can perceive (Geipel et al. 2021) and different innate preferences that determine what stimuli are attractive (Saumweber et al. 2011). Even if two species prefer to associate the same type of cue with food, they might differ in how quickly they can form associations, or how many associations they can learn.

We suggest that future experiments could assess variation in ability to learn cues between bats with different foraging strategies by comparing rates of learning of single cue-types in different modalities. To test the relative use of sound vs odor between species, investigators could repeat this experiment but make the spatial cue unreliable from the start (e.g., Muchhala and Serrano 2015). Our study could only detect the pronounced differences we expected to see between these species; larger sample sizes are necessary to measure subtle differences in cue salience. Due to the logistical difficulties of comparative cognition studies, much clarity would come from greatly increasing the scope and scale of these experiments (e.g., MacLean et al. 2014).

In conclusion, we detected no pronounced difference in cue salience between a bat expected to use primarily odor and spatial cues and a close relative expected to overwhelmingly use acoustic cues. Although there may be differences in cue learning between bat species that we did not detect, our findings show that that larger-scale spatial cues can easily overshadow local features associated with a food source, even in species that feed on mobile prey found in unpredictable locations.

## Acknowledgements

We thank Tate Ackerman, Dylan Valente, Amanda Savage, Dineilys Aparicio, and Vanessa Pérez Pinzón for their assistance with running trials and scoring videos.

## Ethical approval

All experiments were licensed and approved by the Smithsonian Tropical Research Institute (IACUC no. 2017-0102-2020) the Government of Panamá (Ministerio de Ambiente permit SE/A 69-15 and SE/AH-2-6), and by the University of Texas at Austin (AUP-2015-00048).

