## Supplemental materials for "Spatial learning overshadows learning novel odors and sounds in both a predatory and a frugivorous bat"

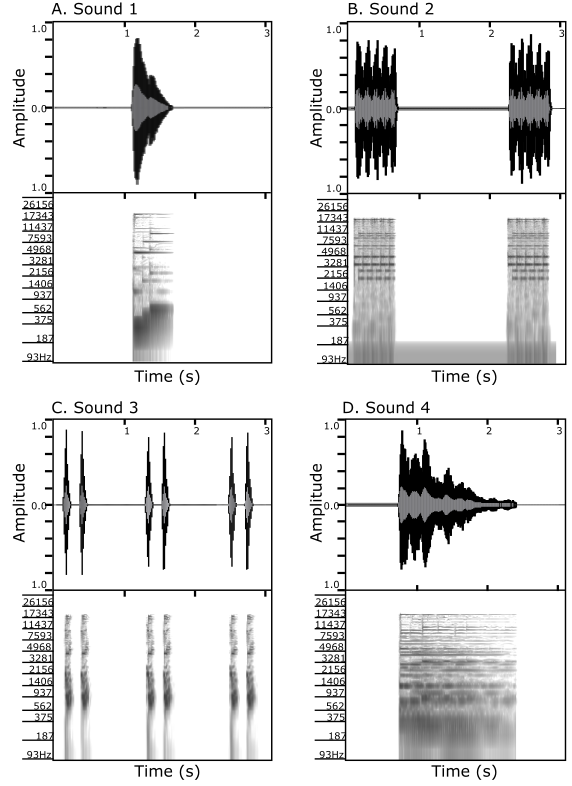

**Figure S1:** Waveforms and spectrograms of the acoustic stimuli used in this experiment.

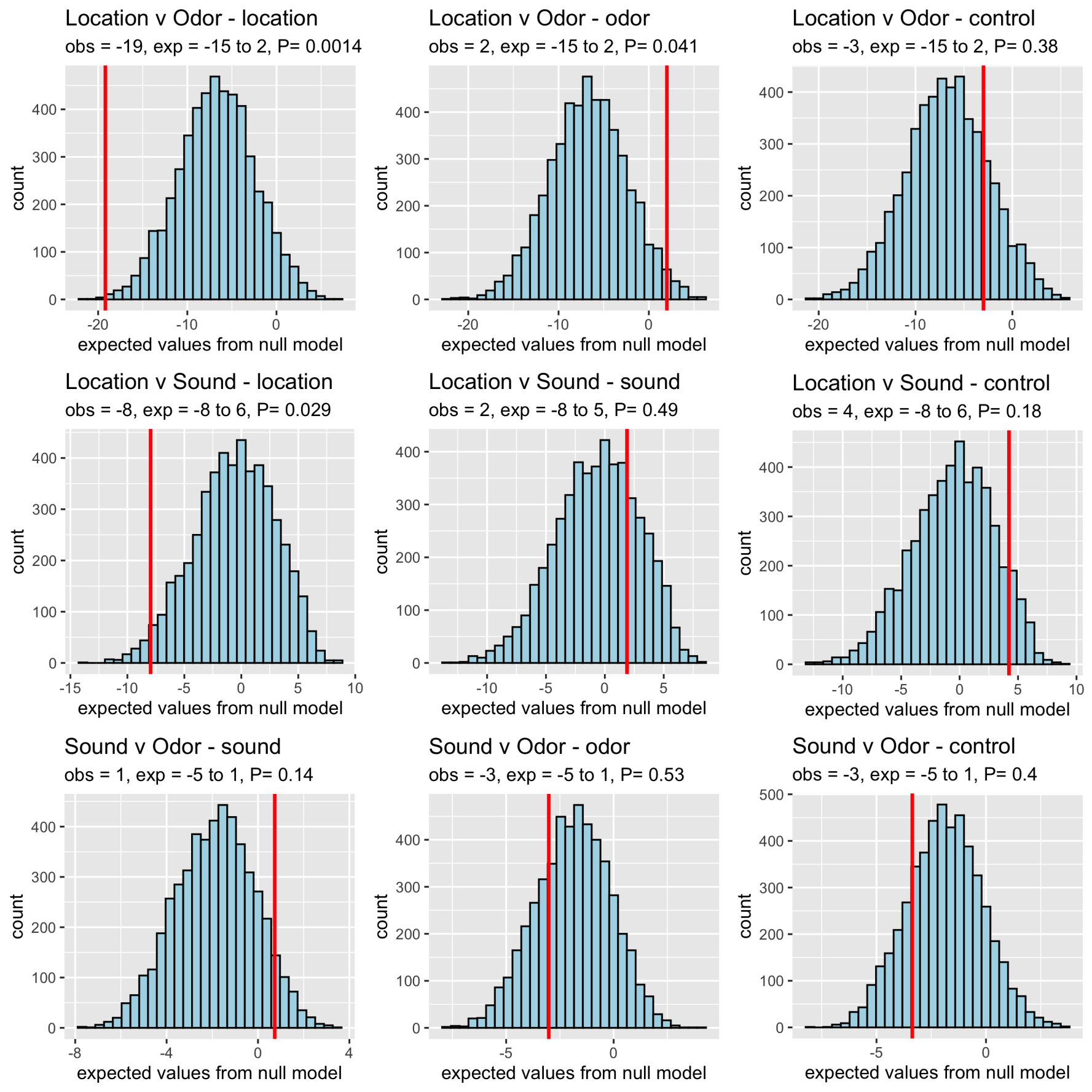
**Figure S2:** Differences in the relative amount that frugivorous *A. jamaicensis* and predatory *L. silvicolum* visited each cue type in the tests. Red lines are the observed species difference (*S*): the mean # visits by *A. jamaicensis* – mean # visits by *L. silvicolum*. Histograms are the null expected difference if the two bats visited the cues randomly, as generated in a permutation test by randomizing the number of visits to each cue within bat (5000 permutations). Two-tailed *P*-values were calculated as the proportion of expected values that had a greater or equal absolute difference to the mean of the expected distribution than *S*. When the line is farther to the left, *L. silvicolum* visited the cue type relatively more than *A. jamaicensis*, and when the line is farther to the right, *A. jamaicensis* visited relatively more.

**Table S1:** Results of previous studies showing which cues animals relied on when given access to multiple cue types.

| **Order** | **Species** | **Context** | **Relevant ecology/ predictions** | **Type of experiment** | **Cues compared** | **Cue(s) relied upon** | **Prediction supported?** | **Citation** |
| --- | --- | --- | --- | --- | --- | --- | --- | --- |
| Aves (Passeriformes) | Black-capped chickadee (*Parus atricapillus*) | Foraging | Caches food- predicted to rely more on spatial cues | Cue dissociations | Spatial (absolute position in a room), spatial (relative position in an array), object (color/pattern) | Spatial  (used object secondarily) | Yes | (Brodbeck 1994) |
|  | Dark-eyed junco (*Junco hyemalis*) | Foraging | Does not cache food- predicted to rely less on spatial cues | Cue dissociations | Spatial (absolute position in a room), spatial (relative positional in an array), object (color/pattern) | Spatial and object equally | Yes | (Brodbeck 1994) |
|  | Marsh tit (*Poecile*  *palustris)* | Foraging | Caches food- predicted to rely more on spatial cues | Cue dissociations | Spatial (positional), color/pattern | Spatial  (used object secondarily) | Yes | (Clayton and Krebs 1994) |
|  | Blue tit (*Cyanistes caeruleus)* | Foraging | Does not cache food- predicted to rely less on spatial cues | Cue dissociations | Spatial (positional), object (color/pattern) | Spatial and object equally | Yes | (Clayton and Krebs 1994) |
|  | Great tit *(Parus major)* **♂** | Foraging | Does not cache food- but predicted males would rely more on spatial cues than females because of trend of larger hippocampi in males | Cue dissociations (foraging in a tray with wells covered with different colors of cloth) | Spatial (position in array), object (color) | Spatial (no evidence of using color cue) | No | (Hodgson and Healy 2005) |
|  | Great tit (*Parus major)* **♀** | Foraging | Does not cache food- but predicted females would rely less on spatial cues than males, b/c of broader trend of smaller hippocampi in females | Cue disassociations (foraging in a tray with wells covered with different colors of cloth) | Spatial (position in array), object (color) | Spatial (no evidence of using color cue) | No | (Hodgson & Healy, 2005) |
|  | Eurasian jay (*Garrulus glandarius*) | Foraging | Caches food- predicted to rely more on spatial memory | Cue dissociations | Spatial (positional), object (color/pattern) | Spatial  (used color/pattern secondarily) | Yes | Clayton and Krebs 1994 |
|  | Jackdaw (*Corvus monedula*) | Foraging | Does not cache food- predicted to rely less on spatial cues | Cue dissociations | Spatial (positional), object (color/pattern) | Spatial and color/pattern equally | Yes | Clayton and Krebs 1994 |
|  | Mountain chickadee  *(Poecile gambeli)* | Foraging | Caches food-predicted to rely on spatial cues | Cue dissociations | Spatial (position in an array), object (color) | Color*, spatial secondarily | No* | (LaDage et al. 2009) |
|  | Zebra finch (*Taeniopygia guttata)* | Foraging | Not stated, but do not cache, so probably expected to use spatial cues relatively less | Cue dissociations | Spatial (positional, absolute place in aviary), object (pattern) | Half of individuals used spatial, half used pattern | Yes | (Mayer and Bischof 2012) |
|  | European greenfinch  *(Chloris chloris)* | Foraging | Are visual foragers that eat ephemeral food- expected to prefer object cues | Cue dissociations- but only rewarded cues were available during training | Spatial (positional), object (color) | With single training trials- color. With multiple training trials, spatial | Yes, contextually | (Herborn et al. 2011) |
| Aves (Galliformes) | Domestic chick (*Gallus gallus domesticus*)  **♂** | Foraging | Visual foragers, males may have territories that prime them to attend to spatial cues | Cue dissociations | Spatial (positional), object (color) | Spatial, color secondarily | NA | (Vallortigara 1996) |
|  | Domestic chick  (*Gallus gallus domesticus*) **♀** | Foraging | Visual foragers | Cue dissociations | Spatial (positional), object (color) | Color | Yes | (Vallortigara 1996) |
| Aves (Apodiformes) | Rufous hummingbird (*Selasphorus rufus*) | Foraging | Feed on a renewable food- predicted to benefit from spatial memories of where that food can be found | Cue dissociations | Spatial, object (color/pattern) | Spatial (color/ pattern secondarily) | Yes | (Hurly and Healy 1996; Hurly and Healy 2002) |
| Aves (Columbiformes) | Homing pigeon (*Colomba livia)* | Homing / foraging | Not specified | Cue dissociations - two experiments | Experiment 1: spatial, object (color)  Experiment 2:  Distal and proximal spatial, object (color) | Experiment 1: spatial    Experiment 2: All three cues (whichever choice had more of the rewarded cues, though spatial was possibly slightly more dominant). | NA | (Strasser and Bingman 1996) |
|  | Homing pigeon (*Colomba livia)* | Foraging | Not specified | Cue dissociations | Spatial (geometric), object (color) | Spatial (geometric), color secondarily | NA | (Vargas et al. 2004; Nardi and Bingman 2007) |
| Mammalia (Primates) | Golden lion tamarin (*Leontopithecus rosalia*) | Foraging | Forage over a relatively wide area to find ripe fruit and insect foraging sites - predicted to rely more on spatial cues | Separate discrimination learning tests - one where animals were trained to find food cued by a location, and a second to find food associated with a color | Spatial, object (color) | Performed better at spatial task. Tamarins performed better at 24 hrs | Somewhat | (Platt et al. 1996) |
|  | Wied's marmoset (*Callithrix kuhli*) | Foraging | Have adaptations to feed on gum, meaning they feed locally on a few trees- predicted to rely less on spatial cues | Separate discrimination learning tests - one where animals were trained to find food cued by a location, and a second to find food associated with a color | Spatial, object (color) | Performed better on spatial task. Marmosets performed relatively better at 5 minutes | Yes | (Platt et al. 1996) |
|  | Orangutan  (*Pongo pygmaeus*) | Foraging | NA | Cue dissociations, discrimination learning | Spatial (position in array), object (color, shape, texture) | Spatial | NA | (Haun et al. 2006) |
|  | Gorilla  *(Gorilla gorilla)* | Foraging | NA | Cue dissociations, discrimination learning | Spatial (position in array), object (color, shape, texture) | Spatial | NA | (Haun et al. 2006) |
|  | Bonobo  *(Pan paniscus)* | Foraging | NA | Cue dissociations, discrimination learning | Spatial (position in array), object (color, shape, texture) | Spatial | NA | (Haun et al. 2006) |
|  | Chimpanzee  *(Pan troglodytes)* | Foraging | NA | Cue dissociations, discrimination learning | Spatial (position in array), object (color, shape, texture) | Spatial | NA | (Haun et al. 2006) |
|  | Human 1-year-old  *(Homo sapiens)* | Received toy | NA | Cue dissociations, discrimination learning | Spatial (position in array), object (color, shape, texture) | Spatial | NA | (Haun et al. 2006) |
|  | Human 3-year-old  *(Homo sapiens)* | Received toy | NA | Cue dissociations, discrimination learning | Spatial (position in array), object (color, shape, texture) | Object | NA | (Haun et al. 2006) |
| Mammalia (Chiroptera) | *Glossophaga commissarisi* | Foraging | Find spatially stable flowers: Feed on a renewable food- predicted to benefit from spatial memories of where that food can be found | Cue dissociations | Spatial (configuration/ global) / shape of flowers (echoacoustic cue) | Spatial (color/pattern secondarily) | Yes | (Thiele and Winter 2005) |
|  | *Glossophaga soricina* | Foraging | Find spatially stable flowers: Feed on a renewable food- predicted to benefit from spatial memories of where that food can be found | Cue dissociations | Spatial (position in array), 2 object cues: odor, echoacoustic shape | Spatial, echoacoustic and odor secondarily | Yes | (Carter et al. 2010) |
|  | *Carollia perspicillata* | Foraging | Feed on fruit- while profitable trees are spatially stable, less need for spatial learning than finding an exact flower | Cue dissociations | Spatial (position in array), 2 object cues: odor, echoacoustic shape | Spatial, echoacoustic and odor secondarily | Somewhat- used spatial but not differently than *G. soricina* | (Carter et al. 2010) |
| Mammalia (Rodentia) | Fox squirrel  *Sciurus niger* | Foraging | Scatter-hoarding- predicted to primarily use distal spatial cues | Cue dissociations | Spatial (positional), spatial (absolute), featural (color/shape) | Spatial cues (absolute), secondarily spatial cues (positional). However, in one condition, the featural cues were used. | Mixed | (Waisman and Jacobs 2008) |
|  | Columbian ground squirrel  *Spermophilus columbianus* | Foraging | Need to locate escape tunnels quickly- no specific prediction | Cue dissociations | Spatial (global landmarks), spatial (local landmarks), spatial (arrangement of array), spatial (route) | Spatial (global landmarks such as trees), used local landmarks (sticks, balls) to some extent | NA | (Vlasak 2006) |
| Reptilia (Squamata) | Whiptail lizard  *(Cnemidophorus inornatus)* | Avoiding heat/ finding shelter | Not specified | Measured relative rates of acquisition of a discrimination task | Spatial (positional), object (color/ pattern/brightness) | Learned spatial faster | NA | (Day et al. 2003) |
| Reptilia (Crocodilia) | Spectacled caiman  *(Caiman crocodilus)* | Avoiding shock | Visual hunters- predicted to use object cues** | Cue dissociation- T-maze, rate of acquisition | Spatial (positional), object (brightness) | Spatial (no evidence of use of color) | No | (Williams 1967a; Williams 1967b) |
| Amphibia (Anura) | Terrestrial toad  *(Rhinella arenarum)* | Finding water | Visual hunters- predicted to use object/ landmark cues** | Cue dissociation- T-maze | Spatial (positional), proximal object cue (color/pattern) | Spatial (second choices not tested) | No | (Daneri et al. 2011) |
|  | Cane toad  *(Rhinella marina)* | Avoiding shock | Visual hunters- predicted to use object/ landmark cues* | Cue dissociation- T-maze, rate of acquisition | Spatial (positional), object (color).  Spatial along | Spatial  (color secondarily- learned confounded cue faster than position alone) | No | (Williams 1967c) |
| Actinopterygii  (Gasterosteiformes) | Three spined stickleback  (*Gasterosteus aculeatus*) - pond population | Foraging | In pond populations, spatial cues are expected to be stable and relatively good predictors for navigation and food finding. | Cue dissociations | Water flow direction / object landmarks (rocks and plants) | The pond population relied more on landmarks to find food than flow direction | Yes | (Girvan and Braithwaite 2000; Braithwaite and Girvan 2003) |
|  | Three spined stickleback  (*Gasterosteus aculeatus*) - river population | Foraging | In river populations, spatial landmark cues are relatively less stable and reliable, but flow direction is more likely to be a navigation cue. | Cue dissociations | Water flow direction / object landmarks (rocks and plants) | The stream population relied relatively more on flow direction than landmarks | Yes | (Girvan and Braithwaite 2000; Braithwaite and Girvan 2003) |

* In this test the spatial task was intentionally designed to see if caching birds *can* use object cues to find food. There were 16 different locations to learn, but only two colors.

**Test was in a different domain than the proposed ecological prediction (e.g.: predicted to use object cues because of foraging strategy, but test did not include foraging).
